# Regulatory Mimicry of Cyclin-Dependent Kinases by Conserved Herpesvirus Protein Kinases

**DOI:** 10.1101/2023.10.18.563025

**Authors:** Naoto Koyanagi, Kowit Hengphasatporn, Akihisa Kato, Moeka Nobe, Kosuke Takeshima, Yuhei Maruzuru, Katsumi Maenaka, Yasuteru Shigeta, Yasushi Kawaguchi

**Author notes:** Correspondence Dr. Yasushi Kawaguchi.

## Abstract

Herpesviruses encode conserved protein kinases (CHPKs) that target cellular cyclin-dependent kinase (CDK) phosphorylation sites; thus, they are termed viral CDK-like kinases. Tyrosine 15 in the GxGxxG motifs of CDK1 and CDK2, whose phosphorylation down-regulates their catalytic activities, is conserved in the corresponding motifs of CHPKs. We found that herpes simplex virus 2 (HSV-2) CHPK UL13 mimicked the regulatory mechanism of CDKs. This regulatory mimicry was conserved in CHPKs encoded by herpesviruses subclassified into subfamilies other than HSV-2, suggesting CHPKs have regulatory and functional mimicry with CDKs. Phosphorylation of the corresponding Tyr in HSV-2 UL13 was required for the down-regulation of viral replication and pathogenicity, specifically in the central nervous system of mice, and for efficient viral recurrence in guinea pigs. These data highlight the dual impact of the regulatory mimicry of CDKs by CHPK on the fine-tuned regulation of lytic and latent HSV-2 infections *in vivo*.

## INTRODUCTION

Viruses require host cellular machinery for proliferation and have evolved multiple strategies to control host cellular machinery to establish a cellular environment favorable for replication and survival. One viral strategy is mimicking key regulatory factors of host cells to control cellular machinery^1–3^. Accumulating evidence suggests viruses encode factors that mimic the functions of various cellular proteins^1–3^. In contrast, regulatory mechanisms of viral factors that mimic cellular factors are likely to be totally different from those of the corresponding cellular proteins. This is because viral factors are expressed from the viral genome and cellular environment are transformed by viral infection where host protein synthesis is shut-off. Furthermore, many cellular pathways in uninfected cells including cell cycle, proliferation, intracellular trafficking, and protein degradation pathways are de-regulated.

Protein phosphorylation, a reversible modification in eukaryotic cells that controls proteins involved in cellular machinery^4–6^, might be a key target for viral hijacking of the host cellular machinery^7,8^. Consistent with this, viral protein kinases conserved in members of the family *Herpesviridae* (herpesviruses), designated conserved herpesvirus protein kinases (CHPKs)^7,9^, share some activities with cellular cyclin-dependent kinases (CDKs); thus, they are also termed viral CDK-like kinases^7,9–18^. CDKs regulate various cellular processes including cell cycle, transcription, metabolism, apoptosis, and proliferation in host cells^19–23^. Like other kinases, each CDK consists of two structurally and functionally distinct lobes (N- and C-lobes). The N-lobe contains a highly conserved GxGxxG motif and phosphorylation of its serine/threonine and tyrosine (Thr-14 and Tyr-15 in CDK1 and -2) inhibits catalytic activity^24^. Of Note, Tyrs in the GxGxxG motifs of CDKs are conserved in the corresponding motifs of CHPKs (S-Fig. 1).

Herpesviruses, double-stranded DNA viruses that are ubiquitous pathogens in mammals, birds, and reptiles^25^, are subdivided into three subfamilies: *Alphaherpesvirinae*, *Betaherpesvirinae,* and *Gammaherpesvirinae*. Herpes simplex virus 2 (HSV-2) in the subfamily *Alphaherpesvirinae*, causes human mucocutaneous and skin diseases (genital herpes, herpetic whitlow), meningitis, and neonatal diseases (life-threatening encephalitis), and is associated with increased risk for human immunodeficiency virus infection^26^. HSV-2 establishes life-long infection in humans with two distinct (lytic and latent) phases of infection. After the initial HSV-2 infection, it becomes latent and frequently reactivates to cause lesions^26^. Here, we focused on HSV-2 CHPK UL13 and investigated whether HSV-2 UL13 mimics the regulatory mechanism of CDKs mediated by Tyr-phosphorylation in the GxGxxG motifs.

## RESULTS

### Tyr phosphorylation in GxGxxG motif of HSV-2 UL13 in infected cells

Tyr at position 162 (Tyr-162) in the GxGxxG motif of HSV-2 UL13, which corresponds to Tyr-15 of CDK1, was well conserved in CHPKs from all three subfamilies examined (S-Fig. 1a, b).

To examine whether UL13 Tyr-162 was phosphorylated in HSV-2-infected cells, human osteosarcoma U2OS cells were infected with wild-type HSV-2 186, a UL13-null mutant virus ΔUL13^27^, UL13-Y162F in which UL13 Tyr-162 was replaced with phenylalanine (Y162F), or its repaired virus UL13-Y162E/F-repair (S-Fig. 2), treated with or without a phosphatase inhibitor, sodium orthovanadate (SOV) from 22 h after infection for an additional 2 h, lysed, and subjected to immunoblotting with anti-UL13-Y162^P^ polyclonal antibodies that specifically react with a peptide corresponding to UL13 residues 157-167 with phosphorylated Tyr-162 (Y162^P^) (S-Fig. 3). In the presence of SOV, anti-UL13-Y162^P^ antibodies reacted with UL13 in lysates of cells infected with wild-type HSV-2 186 or UL13-Y162E/F-repair (Fig. 1a, b). In contrast, anti-UL13-Y162^P^ antibodies barely reacted with UL13 in the lysates of cells mock-infected or infected with ΔUL13 or UL13-Y162F in the presence of SOV. In the absence of SOV, UL13 levels detected by anti-UL13-Y162^P^ antibodies in lysates of cells infected with wild-type HSV-2 186 were very low and required long exposure for immunoblotting (Fig. 1a). Phosphatase treatment of lysates of cells infected with wild-type HSV-2 186 abolished UL13 detection by anti-UL13-Y162^P^ antibodies (Fig. 1c). These results indicated UL13 was phosphorylated at Tyr-162 (UL13-Y162^P^) in HSV-2-infected cells; however, this was unstable and was immediately de-phosphorylated by a phosphatase(s).

**Fig. 1.**
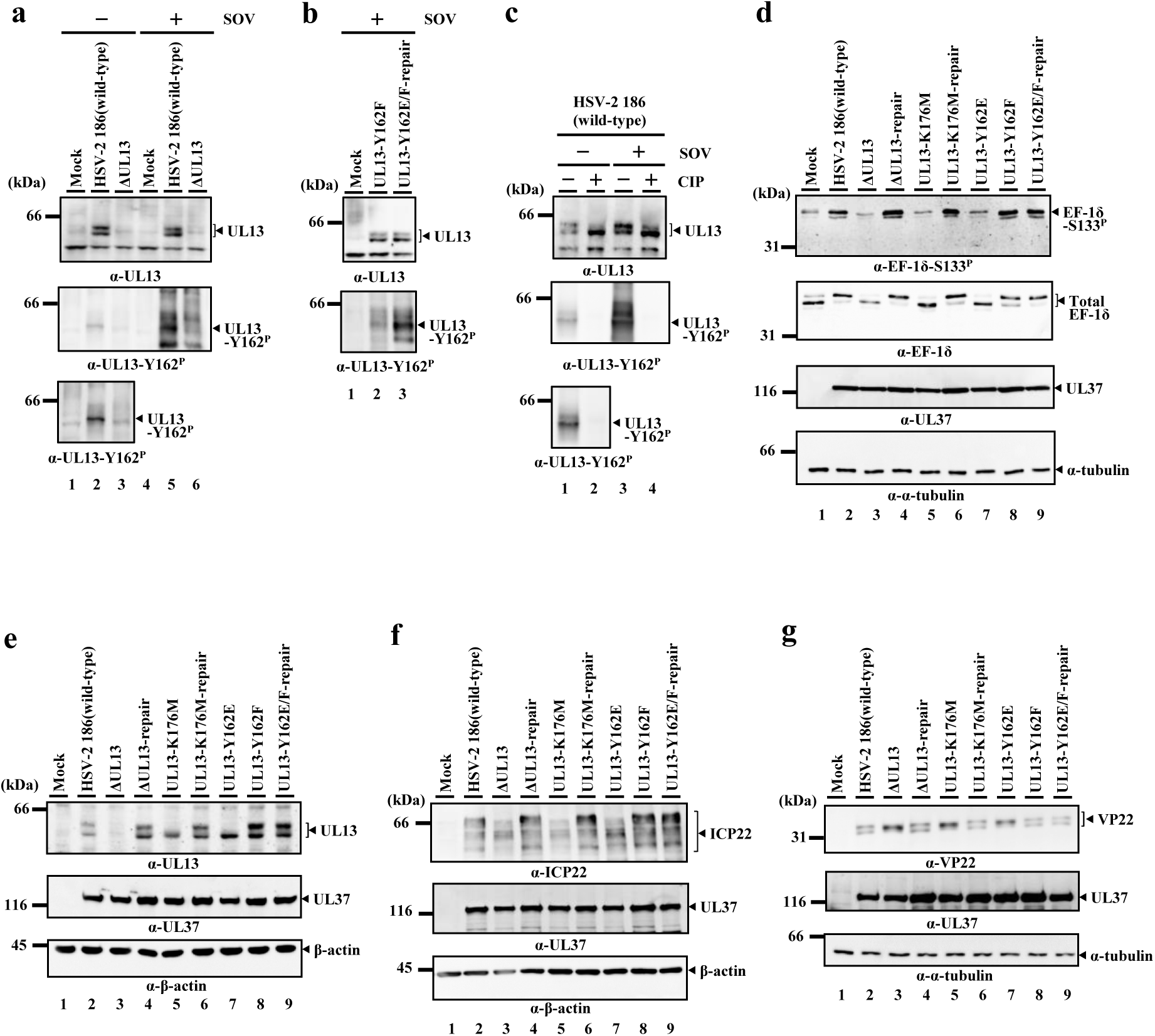
Phosphorylation of UL13 at Tyr-162 and effects of mutations in UL13 on UL13 substrates in HSV-2 infected cells. **a, b.** U2OS cells were mock-infected (a, b) or infected with wild-type HSV-2 186 (a, b), ΔUL13 (a), UL13-Y162F (b), or UL13-Y162E/F-repair (b) at an MOI of 3, incubated with or without 5 mM SOV at 22 h post-infection, harvested at 24 h post-infection, and lysates were analyzed by immunoblotting with antibodies to UL13 or UL13-Y162^P^. **c.** U2OS cells were infected with wild-type HSV-2 186 at an MOI of 3, incubated with or without 5 mM SOV at 22 h post-infection, harvested at 24 h post-infection, lysed, cell lysates were mock-treated or treated with CIP, and then analyzed as described in panel a. **d-g.** U2OS cells mock-infected or infected with wild-type HSV-2 186, ΔUL13, ΔUL13-repair, UL13-K176M, UL13-K176M-repair, UL13-Y162E, UL13-Y162F, or UL13-Y162E/F-repair for 24 h at an MOI of 3 were analyzed by immunoblotting with antibodies to EF-1δ (d), EF-1δ-S133^P^ (d), UL13 (e), ICP22 (f), VP22 (g), UL37 (d-g), α-tubulin (d, g) or β-actin (e, f). Digital images are representative of three independent experiments.

### Effects of HSV-2 UL13 Tyr-162 phosphorylation on UL13 substrates in infected cells

To examine the effects of UL13 phosphorylation at Tyr-162 on UL13 substrates in HSV-2-infected cells, U2OS cells were mock-infected or infected with wild-type HSV-2 186, ΔUL13, ΔUL13-repair^27^, recombinant virus UL13-K176M encoding an enzymatically inactive mutant of UL13^27^, its repaired virus UL13-K176M-repair^27^, UL13-Y162E carrying a phosphomimetic mutation at Tyr-162, UL13-Y162F or UL13-Y162E/F-repair in which mutations in UL13-Y162F and UL13-Y162E were repaired (S-Fig. 2), lysed and subjected to immunoblotting.

Elongation factor 1δ (EF-1δ) is a cellular substrate of CHPKs and CDK1 that phosphorylate Ser-133 in this protein^10^. Infection of cells with wild-type HSV-2 186, ΔUL13-repair, or UL13-K176M-repair increased EF-1δ phosphorylation levels at Ser-133 (EF-1δ-S133^P^) detected by anti-EF-1δ-S133^P^ monoclonal antibody^27^ and the hyper-phosphorylated form of EF-1δ, detected as a slower migrating band by immunoblotting with anti-EF-1δ polyclonal antibodies, compared to mock-infection, but not infection of cells with ΔUL13 or UL13-K176M (Fig. 1d). The increase in the hyper-phosphorylated form of EF-1δ in HSV-2-infected cells resulted from EF-1δ phosphorylation at Ser-133 by UL13^10^. EF-1δ phosphorylation levels at Ser-133 in UL13-Y162E infected cells and the hyper-phosphorylated form of EF-1δ were lower than in cells infected with wild-type HSV-2 186 or UL13-Y162E/F-repair but were similar to those in cells mock-infected or infected with ΔUL13 or UL13-K176M. EF-1δ phosphorylation levels at Ser-133 and the hyper-phosphorylated form of EF-1δ in cells infected with UL13-Y162F was similar to that in cells infected with wild-type HSV-2 186 or UL13-Y162E/F-repair.

Auto-phosphorylated UL13, detected as a slow migrating band by immunoblotting with anti-UL13 polyclonal antibodies^27^, was barely detectable in cells infected with UL13-Y162E, similar to cells infected with UL13-K176M (Fig. 1e). In contrast, auto-phosphorylated UL13 was clearly detected in cells infected with wild-type HSV-2 186, UL13-Y162F, or each repaired virus. Similarly, the phosphorylation status of other UL13 substrates including ICP22 and VP22, detected as differently migrating bands of substrates by immunoblotting^28,29^, in cells infected with UL13-Y162E could not be differentiated from that in cells infected with ΔUL13 or UL13-K176M, but was different from that in cells infected with wild-type HSV-2 186, UL13-Y162F, or each repaired virus (Fig. 1f, g).

Similar results observed in infected U2OS cells were obtained with infected simian kidney epithelial Vero cells (S-Fig. 4). These results indicated the phosphomimetic mutation at HSV-2 UL13 Tyr-162 reduced phosphorylation of all UL13 substrates tested in HSV-2-infected cells to levels comparable with the kinase-dead UL13-K176M mutation, suggesting UL13 phosphorylation at Tyr-162 down-regulated UL13 kinase activity in HSV-2-infected cells.

### Effects of Tyr-phosphorylation in GxGxxG motifs of CHPKs encoded by β- and γ-herpesviruses on EF-1δ phosphorylation

U69 and BGLF4 are CHPKs encoded by a β-herpesvirus (human herpesvirus 6 B [HHV6B]) and γ-herpesvirus (Epstein-Barr virus [EBV]), respectively^30,31^. To examine whether the down-regulation of HSV-2 UL13 by Tyr 162 phosphorylation was conserved in these CHPKs, simian kidney epithelial COS-7 cells were transfected with a plasmid expressing Flag-tagged EF-1δ fused to enhanced green fluorescence protein (EGFP) [EGFP-EF-1δ(F)]^27^ in combination with each of the plasmids expressing wild-type CHPKs and their mutants, and subjected to immunoblotting with the anti-EF-1δ-S133^P^ antibody. Phosphorylation levels of EGFP-EF-1δ(F) in the presence of wild-type HHV6B U69 or EBV BGLF4 increased compared to those in the absence of viral kinases or in the presence of their kinase-dead mutants, verifying these CHPKs phosphorylated EF-1δ at Ser-133 (S-Fig. 5). Phosphorylation levels of EGFP-EF-1δ(F) in the presence of U69-Y207F or BGLF4-Y89F, in which each conserved Tyr in the GxGxxG motifs of viral kinases was substituted with phenylalanine, were comparable to those in the presence of wild-type U69 or BGLF4. In contrast, phosphorylation levels of EGFP-EF-1δ(F) in the presence of U69-Y207E or BGLF4-Y89E, each carrying a phosphomimetic mutation at the conserved Tyr, were significantly lower than those in the presence of wild-type U69 or BGLF4. These results suggested that the down-regulation of CHPK by Tyr-phosphorylation in the GxGxxG motif was conserved in all herpesvirus subfamilies.

### Effects of UL13 Tyr-162 phosphorylation on HSV-2 replication and cell-cell spread

To examine the effects of UL13 phosphorylation at Tyr-162 on HSV-2 replication and cell-cell spread in cell cultures, we analyzed progeny virus yields and plaque sizes in U2OS and Vero cells infected with wild-type HSV-2 186 or each UL13-mutant and repaired virus. Progeny virus yields in U2OS cells infected with UL13-Y162E at a multiplicity of infection (MOI) of 0.01 were significantly lower than in cells infected with wild-type HSV-2 186 or UL13-Y162E/F-repair, but similar in cells infected with ΔUL13 or UL13-K176M (Fig. 2a). Progeny virus yields in U2OS cells infected with UL13-Y162F were similar in cells infected with wild-type HSV-2 186 or UL13-Y162E/F-repair. In contrast, progeny virus yields in U2OS cells infected with each virus at an MOI of 3 were similar (Fig. 2b). Progeny virus yields in Vero cells infected with each virus were comparable at MOIs of 0.01 and 3 (S-Fig. 6). In agreement with the growth properties of these viruses at an MOI of 0.01, UL13-Y162E produced smaller plaques than wild-type HSV-2 186, UL13-Y162F, and UL13-Y162E/F-repair, similar size plaques to ΔUL13 and UL13-K176M in U2OS cells (Fig. 2c), and all viruses produced similar size plaques on Vero cells (S-Fig. 6). These results suggested constitutive phosphorylation at HSV-2 UL13 Tyr-162 reduced HSV-2 replication and cell-cell spread to levels comparable with cells infected with ΔUL13 or UL13-K176M dependent on cell type and MOI.

**Fig. 2.**
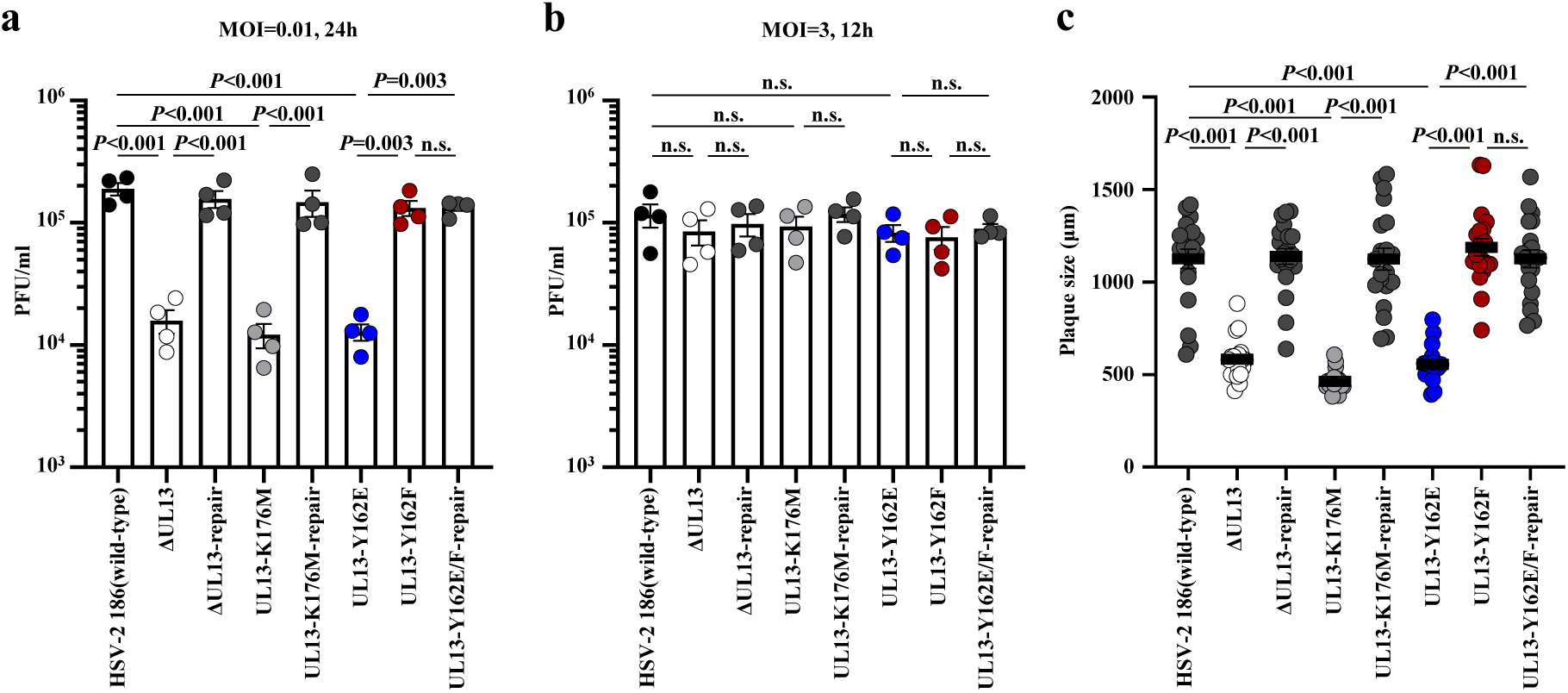
Effects of mutations in UL13 on viral replication and cell-cell spread. **a, b.** U2OS cells were infected with wild-type HSV-2 186, ΔUL13, ΔUL13-repair, UL13-K176M, UL13-K176M-repair, UL13-Y162E, UL13-Y162F, or UL13-Y162E/F-repair at an MOI of 0.01 (a) or 3 (b). Total virus titers in cell culture supernatants and infected cells were harvested at 24 h (a) or 12 h (b) post-infection and assayed. Each value is the mean ± standard error of the mean (SEM) of four experiments. Statistical significance was analyzed by ANOVA with the Tukey’s test. n.s., not significant. **c.** U2OS cells were infected with wild-type HSV-2 186, ΔUL13, ΔUL13-repair, UL13-K176M, UL13-K176M-repair, UL13-Y162E, UL13-Y162F, or UL13-Y162E/F-repair at an MOI of 0.0001 under plaque assay conditions. Diameters of 20 single plaques for each virus were measured at 48 h post-infection. Each data point is the mean ± SEM of the measured plaque sizes. Statistical significance was analyzed by ANOVA with Tukey’s test. n.s., not significant. Data are representative of three independent experiments.

Phenotypes of the Y162F mutation in HSV-2 UL13 suggesting the physiological relevance of UL13 phosphorylation at Tyr-162 were not detected in all cell culture experiments (Fig. 1d-g, Fig. 2, S-Figs. 4, 6), in agreement with the observation that phosphorylation was unstable and UL13-Y162^P^ was immediately de-phosphorylated by a phosphatase(s) in HSV-2-infected cells (Fig. 1a, b).

### Effects of UL13 Tyr-162 phosphorylation on HSV-2 replication and pathogenicity in mice

To examine the effects of UL13 phosphorylation at Tyr-162 on HSV-2 replication and pathogenicity *in vivo*, mice were vaginally infected with UL13-Y162E, UL13-Y162F, or UL13-Y162E/F-repair, and survival, acute genital lesions, and virus titers in vaginal secretions were monitored (Fig. 3a-c). Alternatively, mice were vaginally infected with UL13-Y162E or UL13-Y162E/F-repair, UL13-Y162F or UL13-Y162E/F-repair, or UL13-K176M or UL13-K176M-repair (Fig. 3d-f). At 7 days post-infection, mice were sacrificed, and virus titers in the vagina, spinal cords, and brains were assayed. Survival of mice infected with UL13-Y162E was significantly greater compared with mice infected with UL13-Y162E/F-repair (Fig. 3a). Virus titers in vaginal secretions of mice infected with UL13-Y162E on days 2 and 4, and genital disease scores on days 9 and 12 were significantly lower than in mice infected with UL13-Y162E/F-repair (Fig. 3b, c). Similar survival curves, virus titers in vaginal secretions, and genital disease scores were previously reported with UL13-K176M and UL13-K176M-repair under identical conditions^27^. Virus titers in vagina, spinal cords, and brains of mice infected with UL13-Y162E or UL13-K176M were significantly lower than in mice infected with UL13-Y162E/F-repair or UL13-K176M-repair, respectively (Fig. 3d, e). These results suggested constitutive UL13 phosphorylation at Tyr-162 down-regulated viral pathogenicity and replication in mice during the acute lytic phase of HSV-2 infection.

**Fig. 3.**
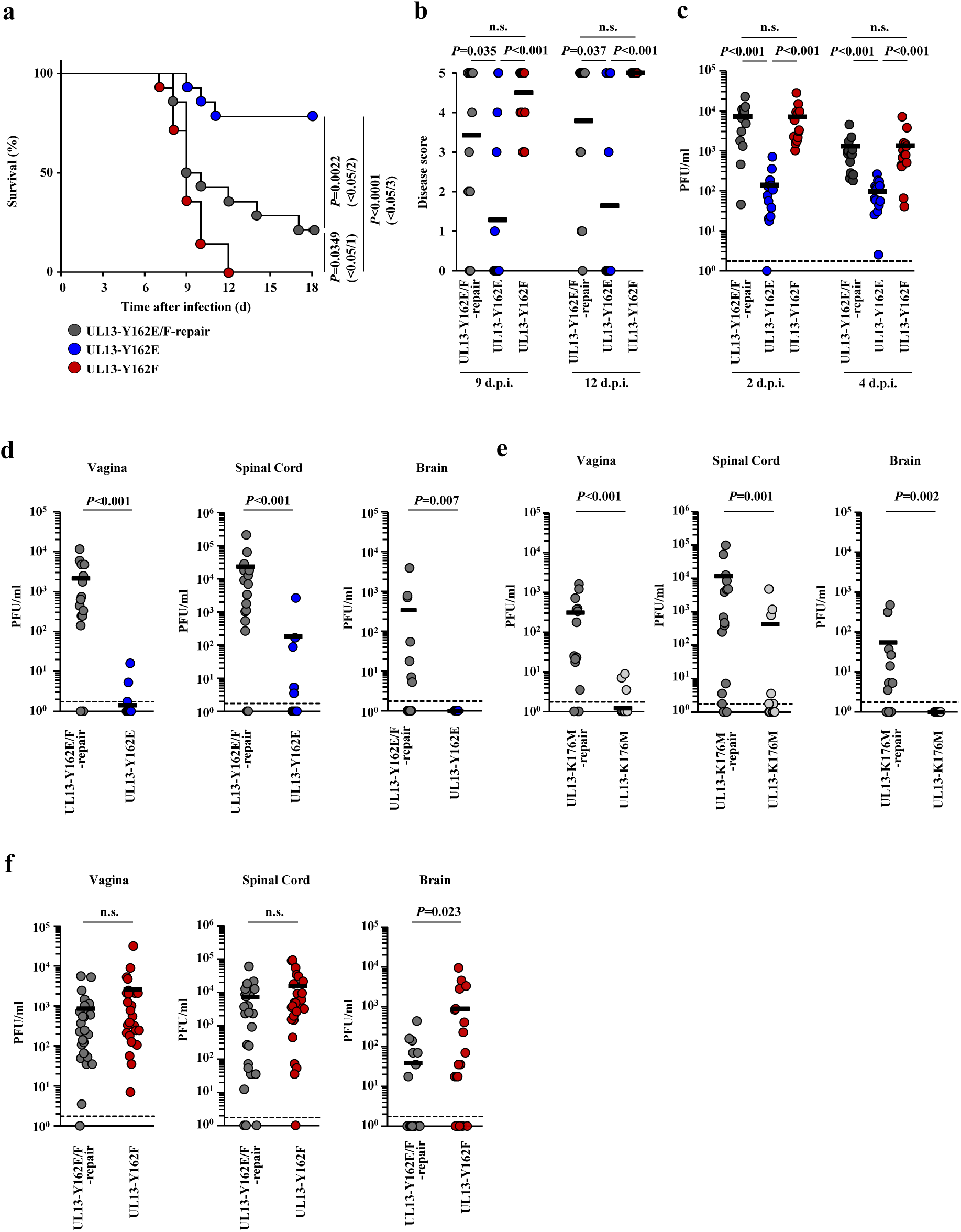
Effects of mutations in UL13 on mortality and viral replication in infected mice following intravaginal infection. **a-c.** Fourteen 6-week-old female ICR mice were pretreated with medroxyprogesterone and the vagina of each mouse was infected with 1×10^4^ PFU UL13-Y162E, UL13-Y162F, or UL13-Y162E/F-repair. (a) Survival of mice was monitored for 18 d post-infection. Differences in the mortality of infected mice were statistically analyzed by the log-rank test, and for three comparison analyses, P values of <0.0167 (0.05/3), <0.025 (0.05/2), or <0.05 (0.05/1) were sequentially considered significant after Holm’s sequentially rejective Bonferroni multiple-comparison adjustment. (b) Clinical scores of infected mice at 9- and 12-days post-infection were monitored. Each data point is the clinical score for one mouse. Horizontal bars indicate the means of each group. Statistical significance values were analyzed by Dunn’s multiple-comparison test. n.s., not significant. (c) Vaginal secretions of infected mice at 2- and 4-days post-infection were harvested, and virus titers were assayed. Each data point is the virus titer in the vaginal secretion of one mouse. Horizontal bars indicate the means of each group. Statistical significance was analyzed by Dunn’s multiple-comparison test. n.s., not significant. The results from three independent experiments were combined. **d-f.** Sixteen (d, e) or 26 (f) 6-week-old female ICR mice were pretreated with medroxyprogesterone and the vaginas of each mouse were infected with 1×10^4^ PFU UL13-Y162E (d) or UL13-Y162E/F-repair (d, f), UL13-K176M (e), UL13-K176M-repair (e), or UL13-Y162F (f). Vaginas, spinal cords, and brains at 7 days post-infection were harvested and virus titers were assayed. Results of three (d, e) or four (f) independent experiments were combined for each virus. Dashed line indicates the limit of detection. Each data point is the virus titer of one mouse. Horizontal bars indicate the mean of each group. Statistical significance was analyzed by Mann-Whitney *U*-test. n.s., not significant.

Whereas UL13-Y162F phenotypes in cell cultures could not be differentiated from those of UL13-Y162E/F-repair (Figs. 1, 2, and S-Figs. 4, 6), the survival of mice infected with UL13-Y162F was significantly lower than mice infected with UL13-Y162E/F-repair (Fig. 3a) and virus titers in brains of mice infected with UL13-Y162F were significantly higher than in mice infected with UL13-Y162E/F-repair (Fig. 3f). In contrast, virus titers in the vagina and spinal cords of mice infected with UL13-Y162F were similar to those in mice infected with UL13-Y162E/F-repair (Fig. 3f). Consistently, acute genital disease scores and virus titers in the vaginal secretions of mice infected with UL13-Y162F were similar to mice infected with UL13-Y162E/F-repair (Fig. 3b, c). These results suggested UL13 phosphorylation at Tyr-162 was required for the down-regulation of viral replication and pathogenicity specifically in the central nervous system (CNS) of mice during the acute lytic phase of infection. In contrast, UL13 phosphorylation at Tyr-162 was not required for viral replication or pathogenic manifestations in the vagina and spinal cords of mice.

### Effects of UL13 Tyr-162 phosphorylation on HSV-2 latency in guinea pigs

To examine the physiological effects of UL13 phosphorylation at Tyr-162 on HSV-2 latency and reactivation, guinea pigs were vaginally infected with UL13-Y162F or UL13-Y162E/F-repair, and survival, acute genital lesions, and virus titers in vaginal secretions were monitored. Survival, acute genital disease scores, and virus titers in vaginal secretions in guinea pigs infected with UL13-Y162F were similar to those infected with UL13-Y162E/F-repair, indicating UL13 phosphorylation at Tyr-162 was not required for viral replication and pathogenic manifestations in the vagina and pathogenicity in guinea pigs (S-Fig. 7). These results confirmed viral replication and pathogenic manifestations in peripheral sites of mice infected with each virus (Fig. 3b-f), but not with the survival of infected mice (Fig. 3a). This difference may be due to guinea pig experiments where HSV-2 infection had lower viral pathogenicity in the CNS, as most (77%) guinea pigs infected with UL13-Y162E/F-repair survived (S-Fig. 7a) unlike the mouse experiments where most (79%) mice infected died (Fig. 3a).

Following recovery from acute infection, evaluable guinea pigs were monitored daily between days 21-56 after infection for recurrent diseases. Guinea pigs infected with UL13-Y162F during the latent phase of infection had significantly reduced mean cumulative recurrence and number of recurrent lesion days compared with guinea pigs infected with UL13-Y162E/F-repair (Fig. 4a, b). After guinea pigs vaginally infected with UL13-Y162F or UL13-Y162E/F-repair recovered from acute infection, these evaluable were sacrificed 21 days after infection and latent HSV-2 genomes in the dorsal root ganglia (DRGs) were quantitated. HSV-2 DNA levels in DRGs during the latent phase of infection in guinea pigs infected with UL13-Y162F were similar to those infected with UL13-Y162E/F-repair (Fig. 4c). Thus, UL13 phosphorylation at Tyr-162 was required for efficient reactivation from latency in guinea pigs although it had no effect on establishing viral latency.

**Fig. 4.**
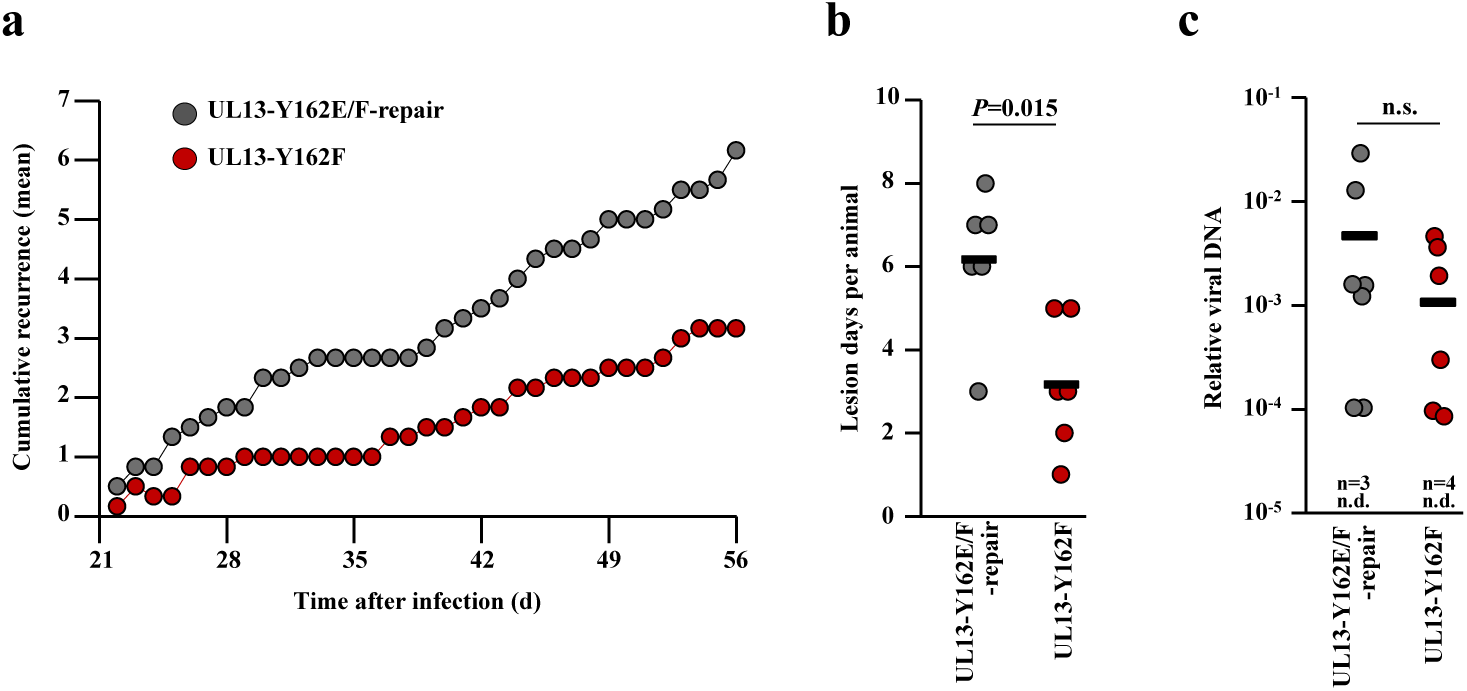
Effects of mutations in UL13 Tyr-162 on HSV-2 latency and recurrence in guinea pigs following intravaginal infection. **a.** Eighteen 5-week-old female Hartley guinea pigs were intravaginally infected with 1×10^4^ PFU UL13-Y162F or UL13-Y162E/F-repair. Guinea pigs with no infectious virus detected in vaginal washes at 1, 3, and 5 days after infection, with no detectable disease by 21 days after infection, with vaginal lesions that had not healed by 21 days after infection, or were dead by 21 days after infection were removed from the analysis, resulting in 6 guinea pigs for each UL13-Y162F and UL13-Y162E/F-repair. Mean number of cumulative recurrences per guinea pig in each group from 22-56 days after infection. Results from two independent experiments were combined. **b.** The number of days with a recurrent lesion is shown in a. Each data point is a recurrent number of one guinea pig. Horizontal bars indicate the mean of each group. Statistical significance was analyzed by Mann-Whitney *U*-test. **c.** Twelve 5-week-old female Hartley guinea pigs were intravaginally infected with 1×10^4^ PFU UL13-Y162F or UL13-Y162E/F-repair. Guinea pigs with no detectable disease or death by 21 days after infection were removed from the analysis, resulting in 10 guinea pigs in the UL13-Y162F and UL13-Y162E/F-repair groups. Twenty-one days after infection, viral genomes from DRG of infected guinea pigs were quantified by ddPCR. Results from two independent experiments were combined. Each data point is the relative amount of each viral genome in the DRG of one guinea pig. Horizontal bars indicate the mean of each group. Statistical significance was analyzed by the Mann-Whitney *U*-test. n.s., not significant. n.d., number of animals with no viral genomes detected in the tissue.

## DISCUSSION

Tyr-15 in the GxGxxG motif of CDK1 and CDK2, whose phosphorylation down-regulates their catalytic activities^32–34^, is well-conserved in most (85%) kinases in the CDK family as observed in CHPKs (S-Fig. 1). Few kinases (11.7%) in other families contain a corresponding Tyr in the GxGxxG motif (S-Fig. 8) and the functional effects of its phosphorylation are unclear^35,36^. The current study showed Tyr phosphorylation in GxGxxG motifs of CHPKs down-regulated their kinase activities indicating CHPKs specifically mimic the regulatory mechanism mediated by Tyr-15 phosphorylation in CDK1 and CDK2. Thus, herpesviruses have evolved functional and regulatory mimicry of CDKs by CHPKs. Interestingly, the corresponding Tyrs in the GxGxxG motif of CDKs and CHPKs are also well conserved in F10L homologs, viral kinases encoded by poxviruses, but not in other viral kinases including B1R homologs of poxviruses and Us3 homologs of alphaherpesviruses^7^ (S-Fig. 9), suggesting poxviruses might have also evolved CDK regulatory mimicry by conserved F10L kinases.

We reported the phenotype of the phosphorylation-null mutation at Tyr-162 (the Y162F mutation) in HSV-2 UL13, suggesting the physiological relevance of the regulatory mimicry of CDKs by HSV-2 UL13, was evident specific occasions *in vivo*. Whereas the phosphorylation-null mutation aberrantly augmented virulence and viral replication in the CNS of mice, but not the vagina or spinal cord during the acute lytic phase of infection, the mutation impaired the recurrence rate in guinea pig vaginas during the latent phase of infection without affecting latent infection of the DRGs. Thus, the regulatory mimicry of CDKs by HSV-2 UL13 seems to be critical not only for the down-regulation of viral replication and pathogenicity specifically in the CNS during the acute lytic phase of infection but also for efficient reactivation from latency. Consequently, the regulatory mimicry of CDKs by HSV-2 UL13 might affect the fine-tuned regulation of acute lytic and latent HSV-2 infections *in vivo*. Herpesviruses can successfully persist over a lifetime and be transmitted to new hosts without causing significant damage, suggesting they have evolved a sophisticated balance with their hosts during a long history of coevloution^37,38^. The negative regulation of HSV-2 UL13 by Tyr-162 phosphorylation might be a viral strategy to co-exist with a host by preventing high CNS pathogenicity during the acute lytic phase of infection, which allows host survival and viral persistence. During reactivation from latency, UL13 might counteract signaling pathway(s) required for efficient reactivation; thus, the phosphorylation-dead mutation at UL13 Tyr-162 that precludes the negative regulation of UL13 activity, might impair reactivation from latency. UL13 homologs inhibit various cellular signaling pathways including JAK/STAT, RIG-I-like receptor, and cGAS/STING^39–42^. It would be of interest to investigate these pathways regulate reactivation from latency. Alternatively, UL13 might promote signaling pathway(s) required for efficient reactivation as a UL13 homolog promoted escape from viral genome silencing in neurons and axonal anterograde transport upon reactivation^43,44^. Outcomes of signaling pathways can depend on fine-tuned activities of kinases that modulate pathways and failure of proper kinase regulation leads to different signaling pathway outcomes^45^. UL13 dysregulation by the phosphorylation-null mutation, which might preclude the fine-tuned down-regulation of UL13 activity, might impair reactivation from latency.

## METHODS

### Cells and viruses

Simian kidney epithelial Vero and COS-7 cells, rabbit skin cells and human osteosarcoma U2OS cells, and HSV-2 wild-type strain HSV-2 186 were described previously^10,46–49^. Recombinant virus HSV-2 ΔUL13 in which the UL13 gene was disrupted by deleting UL13 codons 159-417, recombinant virus HSV-2 ΔUL13-repair in which the UL13 null mutation was repaired, recombinant virus HSV-2 UL13-K176M encoding an enzymatically inactive UL13 mutant in which lysine at UL13 residue 176 was replaced with methionine, and recombinant virus HSV-2 UL13-K176M-repair in which the UL13 K176M mutation was repaired were described previously^27^ (S-Fig. 2).

### Plasmids

pGEX-ICP22-P1 was constructed by amplifying the domains of HSV-2 ICP22 (encoded by ICP22 codons 1–165) from pYEbac861 by PCR using the primers listed in S-Table 1, and cloning it into the *Eco*RI and *Sal*I sites of pGEX-4T-1(GE Healthcare) in frame with glutathione S-transferase (GST) sequences. pME-BGLF4 and pME-BGLF4-K102I were described previously^50^. pME-BGLF4-Y89E or pME-BGLF4-Y89F, in which Tyr-89 of BGLF4 was replaced with glutamic acid or phenylalanine, respectively, were generated according to the manufacturer’s instructions using the QuikChange site-directed mutagenesis XL kit with complementary oligonucleotides listed in S-Table 1, containing a specific nucleotide substitution (Stratagene) based on pME-BGLF4. pcDNA-SE-U69 was constructed by amplifying the entire coding sequence of HHV-6B U69 from HHV-6B strain HST DNA by PCR using the primers listed in S-Table 1, and cloning it into the *Eco*RI and *Not*I sites of pcDNA-SE^51^ in frame with a Strep-tag sequence. pcDNA-SE-U69-K219M, in which Lys-219 of U69 was replaced with methionine, was generated according to the manufacturer’s instructions using the QuikChange site-directed mutagenesis XL kit with complementary oligonucleotides listed in S-Table 1, containing the specific nucleotide substitution (Stratagene) based on pcDNA-SE-U69. pcDNA-SE-U69-Y207E and pcDNA-SE-U69-Y207F, in which Tyr-207 of U69 was replaced with glutamic acid or phenylalanine, respectively, were generated according to the manufacturer’s instructions using the QuikChange site-directed mutagenesis XL kit with complementary oligonucleotides shown in S-Table 1, containing the specific nucleotide substitution (Stratagene) based on pcDNA-SE-U69. pEGFP-EF-1δ(F), in which EF-1δ was tagged with the Flag epitope and EGFP, and pEGFP-EF-1δ-S133A(F), in which EF-1δ Ser-133 was replaced with alanine were described previously^27^.

### Construction of recombinant viruses

Recombinant virus UL13-Y162E, in which Tyr-162 of HSV-2 UL13 was substituted with glutamic acid (S-Fig. 2), was generated by the two-step Red-mediated mutagenesis procedure using *Escherichia coli* (*E. coli*) GS1783 strain containing pYEbac861^27^ as described previously^52,53^, except using the primers listed in S-Table 2. Recombinant virus UL13-Y162F, in which Tyr-162 of UL13 was substituted with phenylalanine (S-Fig. 2), was generated by the two-step Red-mediated mutagenesis procedure using *E. coli* GS1783 strain containing the UL13-Y162E genomes, except using the primers listed in S-Table 2. Recombinant virus UL13-Y162E/F-repair, in which the Y162F mutation in UL13 was repaired (S-Fig. 2), was generated by the two-step Red-mediated mutagenesis procedure, using *E. coli* GS1783 containing the UL13-Y162F genomes, and the primers listed in S-Table 2. UL13-Y162E/F-repair is the repaired virus of UL13-Y162E and UL13-Y162F.

### Production and purification of GST fusion proteins

GST-ICP22-P1 was expressed in *E. coli* Rosetta (Novagen), transformed with pGEX-ICP22-P1, purified by glutathione-sepharose beads (GE Healthcare Life Science), and eluted with GST elution-buffer (50 mM Tris-HCl [pH 8.0], 10 mM reduced glutathione (Sigma)) as described previously^51^.

### Antibodies

Antibodies were as follows: commercial mouse monoclonal antibodies to Flag-tag (M2; Sigma), Strep-tag (4F1; MBL), β-actin (AC15; Sigma), α-tubulin (DM1A; Sigma), and rabbit polyclonal antibodies to UL37 (CAC-CT-HSV-UL37; CosmoBio). Mouse monoclonal antibodies to UL13 and EF-1δ with phosphorylated Ser-133 and rabbit polyclonal antibodies to EF-1δ, BGLF4, and VP22 were reported previously^10,54–57^. Rabbit polyclonal antibodies that recognize UL13 with phosphorylated Tyr-162 was generated by SCRUM Inc. (Tokyo, Japan). As the antigen, the phosphopeptide GGSGG(pY)GEVQL, which corresponds to the UL13 residues 157-167, was synthesized and conjugated at the amino terminus by an additional cysteine to the keyhole limpet hemocyanine. Two rabbits were immunized four times with the antigen mixed with the Freund’s complete adjuvant. The serum from one of the rabbits was subjected to affinity purification using a column conjugated with the UL13 phosphopeptide. The bound antibodies were eluted from the column and passed through another column conjugated with UL13 unphosphorylated peptide Cys-GGSGGYGEVQL to eliminate the antibodies that bound to the unphosphorylated peptide. To generate mouse polyclonal antibodies to HSV-2 ICP22, BALB/c mice were immunized once with purified MBP-ICP22-P1 and TiterMax Gold (TiterMax USA, Inc.). Sera from immunized mice were used as sources of mouse polyclonal antibodies to ICP22.

### ELISA

The specificity of anti-UL13-Y162^P^ polyclonal antibodies was analyzed by enzyme-linked immunosorbent assay (ELISA). Nunc-Immuno plates (Thermo Scientific) coated with the phosphorylated Tyr-162 peptide of UL13 (Cys-GGSGG(pY)GEVQL) or the unphosphorylated Tyr-162 peptide of UL13 (Cys-GGSGGYGEVQL) were blocked with 2% fetal calf serum (FCS) in phosphate-buffered saline (PBS) and anti-UL13-Y162^P^ polyclonal antibodies diluted with 2% FCS in PBS were added to the plates. Anti-rabbit IgG, HRP-linked F (ab’)2 Fragment (GE Healthcare Bio-Sciences) and 1-Step™ TMB ELISA Substrate Solutions (Thermo Scientific) were added to the plates and detected by a Perkin Elmer EnSpire multimode plate reader.

### Immunoblotting

Immunoblotting was performed as described previously^58^. To detect UL13 phosphorylation at Tyr-162, cell lysates in sodium dodecyl sulfate (SDS) sample buffer B (62.5 mM Tris-HCl [pH 6.8], 2% SDS, 20% glycerol, 5% 2-mercaptoethanol, containing protease and phosphatase inhibitor cocktails (Nacalai Tesque)), were used. Brightness/contrast of raw blots were equally adjusted across the entire image with Image lab software (BioRad) to generate representative images. Protein (EGFP-EF-1δ-S133^P^(F)) levels present in immunoblot bands were quantified using the ImageQuant LAS 4000 system with ImageQuant TL7.0 analysis software (GE Healthcare Life Sciences) according to the manufacturer’s instructions and normalized to that of EGFP-EF-1δ(F) proteins and then to the sum of the data across multiple experiments in the same blot, as described previously^59^.

### Inhibitor treatment

U2OS cells infected with wild-type HSV-2 186 or each recombinant virus were treated with or without 5 mM SOV (Wako) at 22 h post-infection for further analyses.

### Phosphatase treatment

Lysates of U2OS cells infected with wild-type HSV-2 186 at an MOI of 3 for 24 h were treated with calf intestinal alkaline phosphatase (CIP) (New England BioLabs) as described previously^60^.

### Determination of plaque size

Vero and U2OS cells were infected with each recombinant virus at an MOI of 0.0001, and plaque sizes were determined as described previously^61^.

### Intravaginal infection of mice

Female ICR mice were purchased from Charles River. For intravaginal HSV-2 infection, 5-week-old ICR mice were injected subcutaneously in the neck ruff with 1.67 mg medroxyprogesterone (Depo-Gestin; A.N.B Laboratories) in 200 μl PBS 7 days prior to viral infection. Treated mice were then infected intravaginally with 1×10^4^ plaque forming unit (PFU) of each virus as described previously^62^. Mice were monitored daily until 18 days post-infection for survival and the severity of vaginal disease using a scoring system as described previously^62^. Virus titers in vaginal secretions of mice were determined as described previously^62^. To determine virus titers in vaginas, spinal cords, and brains of mice, the infected mice at 7 days post-infection were sacrificed, and the tissues were removed, sonicated in 1 ml of medium 199 containing 1% FCS and antibiotics, and frozen at −80°C. Frozen samples were later thawed, and virus titers in the supernatants obtained after centrifugation of the samples were determined by standard plaque assays on Vero cells. All animal experiments were performed in accordance with the Guidelines for Proper Conduct of Animal Experiments, Science Council of Japan. The protocol was approved by the Institutional Animal Care and Use Committee (IACUC) of the Institute of Medical Science, The University of Tokyo (IACUC protocol approval PA11-81, PA16-69, A21-55).

### Intravaginal infection of guinea pigs

Female Hartley strain guinea pigs were purchased from Japan SLC, Inc. For intravaginal infection, 5-week-old female guinea pigs were infected with 1×10^4^ PFU UL13-Y162F or UL13-Y162E/F-repair per vagina. Guinea pigs were monitored daily until 21 days post-infection for survival and the severity of vaginal disease using a scoring system of 0 for no sign of disease, 1 for redness/swelling, 2 for 1-2 lesions, 3 for 3-5 lesions, 4 for ≥6 lesions, the coalescence of lesions, ulcerated lesions, or neurological symptoms, and 5 for death as described previously^63^. Guinea pigs were euthanized after showing signs of severe disease. Vaginal washes of guinea pigs were collected by pipetting 300 µl of medium 199 containing 1% FCS and antibiotics in and out of the vagina 10 times, and diluted to a final volume of 1 ml in medium 199 containing 1% FCS and antibiotics. Virus titers were determined by standard plaque assay. After recovery from acute genital HSV-2 infections, guinea pigs were observed daily from 22-56 days post-infection for recurrent lesions and were assigned a score of 1 point for each day that a lesion was present. Guinea pigs with no infectious virus detected in vaginal washes 5 days post-infection, with no detectable disease by 21 days post-infection, or with vaginal lesions that had not healed by 21 days post-infection were removed from the analysis of HSV-2 recurrence. The protocol was approved by the IACUC of the Institute of Medical Science, The University of Tokyo (IACUC protocol approval PA15-15, A19-91).

### Detection of viral DNA by Droplet Digital PCR

Five-week-old female guinea pigs were intravaginally infected with 1×10^4^ PFU UL13-Y162F or UL13-Y162E/F-repair per vagina. Guinea pigs with no detectable disease or death by 21 days post-infection were removed from the analysis. Total DNA was isolated from DRGs of guinea pigs sacrificed at 21 days post-infection. Total DNA in DRGs was isolated by a GeneJET Genomic DNA Purification Kit (Thermo Fisher Scientific) according to the manufacturer’s instructions. Droplet Digital PCR (ddPCR) was performed to measure HSV-2 genomic DNA levels in DRGs using the QX100 droplet digital PCR system (Bio-Rad Laboratories). HSV-2 genomic DNA was quantified using the following gD primers/TaqMan probe set: 5’-GGTGAAGCGTGTTTACCACA-3’, 5’-TACACAGTGATCGGGATGCT-3’, and a fluorescein amidite (FAM) labeled Universal Probe Library probe 65 (Roche). Cellular genomic DNA was quantified by a RPP30 hexachloro-fluorescein (HEX) assay (BioRad Assay ID: dCNS675240177). The ddPCR reaction mixture consisted of 10 µl ddPCR Supermix for Probe (no dUTP) (Bio-Rad), 0.18 µl each 100 µM HSV-2 gD primer, 0.5 µl Universal Probe Library probe 65 (Roche), 1 µl ddPCR Copy Number Assay for guinea pig RPP30, and 3 µl template DNA in a final volume of 20 µl. Each reaction was mixed with 70 µl Droplet Generation Oil for Probes (Bio-Rad) and loaded into a DG8 cartridge (Bio-Rad). A QX200 Droplet Generator (Bio-Rad) was used to make the droplets, which were transferred to a 96-well plate and the following PCR reaction was run: 95°C for 10 minutes, 40 cycles of 94°C for 30 seconds, and 54°C for 1 minute, followed by 98°C for 10 minutes, ending at 4°C. The ramp rate was 2℃/sec for all steps. The QX200 Droplet Reader (Bio-Rad) was used to analyze droplets for fluorescence measurement of the FAM and HEX probes. Data were analyzed in QX Manager 1.1 Standard Edition (Bio-Rad). To determine the relative viral genomic DNA levels, the number of HSV-2 gD-positive droplets was divided by the number of RPP30-positive droplets in the same 20 μl reaction.

### Statistical analysis

Differences in viral replication and plaque size in cell cultures, and relative amounts of phosphorylated EF-1δ were analyzed statistically by analysis of variance (ANOVA) followed by Tukey’s post-hoc test. Differences in viral replication in vaginas, spinal cords, and brains of mice, viral replication in vaginal washes, disease scores, relative amount of latent HSV-2 genome, mean number of recurrences in guinea pigs, and ELISA results were statistically analyzed by the Mann–Whitney *U-*test. Differences in viral replication in vaginal washes and disease scores of mice were analyzed statistically by Dunn’s multiple comparisons test. Differences in the mortality of infected guinea pigs were statistically analyzed by the log-rank test. A P value of <0.05 was considered statistically significant. Differences in the mortality of infected mice were statistically analyzed by the log-rank test, and for the three comparison analyses, P values of <0.0167 (0.05/3), <0.025 (0.05/2), or <0.05 (0.05/1) were sequentially considered significant after Holm’s sequentially rejective Bonferroni multiple-comparison adjustment. All statistical analyses were performed with GraphPad Prism 8 (GraphPad Software, San Diego, CA).

## Supporting information

S-Figure ans S-Table

## ACKNOWLEDGEMENTS

We thank Risa Abe, Tohru Ikegami, Yui Muto, Keiko Sato and Yoshie Asakura for their excellent technical assistance. We are grateful to Yasuko Mori and Jun Arii for providing valuable reagents, and Keizo Tomonaga and Junna Kawasaki for helpful discussions. This study was supported by Grants for Scientific Research and Grant-in-Aid for Scientific Research (S) (20H05692) from the Japan Society for the Promotion of Science (JSPS), grants for Scientific Research on Innovative Areas (21H00338, 21H00417, 22H04803) and a grant for Transformative Research Areas (22H05584) from the Ministry of Education, Culture, Science, Sports and Technology of Japan, a PRESTO grant (JPMJPR22R5) from Japan Science and Technology Agency (JST), grants (JP20wm0125002, JP22fk0108640, JP22gm1610008, JP223fa627001, JP23wm0225031, JP23wm0225035) from the Japan Agency for Medical Research and Development (AMED), grants from the International Joint Research Project of the Institute of Medical Science, the University of Tokyo, grants from the Takeda Science Foundation, the Mitsubishi Foundation, the Uehara Memorial Foundation, and the Waksman Foundation of Japan, and the GSK Japan Research Grant 2019.

**S-Fig. 1. Sequence alignment around the GxGxxG motif of CHPKs and human CDKs. a.** CHPKs amino acids are labeled with their NCBI gene identification numbers and virus names. HSV-1, herpes simplex virus 1; HSV-2, herpes simplex virus 2; VZV, varicella-zoster virus; PRV, pseudorabies virus; MDV, Marek’s disease virus; SaHV-1, saimiriine herpesvirus 1; CaHV-1, canid herpesvirus 1; FeHV-1, feline herpesvirus 1; BoHV-1, bovine herpesvirus 1; CpHV-1, caprine herpesvirus 1; EHV-1, equine herpesvirus 1; EHV-4, equine herpesvirus 4; HCMV, human cytomegalovirus; HHV-6A, human herpesvirus 6A; HHV-6B, human herpesvirus 6B; HHV-7, human herpesvirus 7; MCMV, murine cytomegalovirus; RhCMV, rhesus cytomegalovirus; EBV, Epstein-Barr virus; KSHV, Kaposi’s sarcoma-associated herpesvirus; MHV-68, murine gammaherpesvirus-68; EHV-2, equine herpesvirus 2; and OvHV-2, ovine herpesvirus 2. HSV-1, HSV-2, VZV, PRV, MDV, SaHV-1, CaHV-1, FeHV-1, BoHV-1, CpHV-1, EHV-1 and EHV-4 belong to the subfamily *Alphaherpesvirinae*; HCMV, HHV-6A, HHV-6B, HHV-7, MCMV and RhCMV belong to the *Betaherpesvirinae*; and EBV, KSHV, MHV-68, EHV-2 and OvHV-2 belong to the *Gammaherpesvirinae*. Highly conserved glycine residues in the GxGxxG motif and valine residue near the GxGxxG motif are in white. Tyrosine residues corresponding to the CDK1 Tyr-15 position of the GxGxxG motif are in pink. **b.** CDK amino acids are labeled with their NCBI gene identification numbers. Highly conserved glycine residues in the GxGxxG motif and valine residue near the GxGxxG motif are in white. Tyrosine residues corresponding to the CDK1 Tyr-15 position of the GxGxxG motif are in pink.

**S-Fig. 2. Schematic diagrams of the genomic structures of wild-type HSV-2 186 and the relevant domains of recombinant viruses used in this study.** Line 1, wild-type HSV-2 186 genome; Line 2, domain of the UL12 gene to the UL15 gene; Lines 3 to 10, recombinant viruses with mutations in the UL13 gene.

**S-Fig. 3. Generation of rabbit polyclonal antibodies to UL13-Y162^P^**. ELISA was performed to assess the specificity of UL13-Y162^P^ polyclonal antibodies. Phosphorylated Tyr-162 peptide of UL13 (Cys-GGSGG(pY)GEVQL) or non-phosphorylated Tyr-162 peptide of UL13 (Cys-GGSGGYGEVQL) was used for ELISAs. Each value is the mean ± SEM of four experiments. Statistical significance was analyzed by Mann-Whitney *U*-test.

**S-Fig. 4. Effects of mutations in UL13 on post-translational processing of UL13 substrates in Vero cells.** Vero cells mock-infected or infected with wild-type HSV-2 186, ΔUL13, ΔUL13-repair, UL13-K176M, UL13-K176M-repair, UL13-Y162E, UL13-Y162F, or UL13-Y162E/F-repair for 24 h at an MOI of 3 were analyzed by immunoblotting with antibodies to EF-1δ (a), EF-1δ-S133^P^ (a), UL13 (b), ICP22 (c), VP22 (d), UL37 (a-d), α-tubulin (a, b, d), or β-actin (c). Digital images are representative of three independent experiments.

**S-Fig. 5. Effects of mutations in tyrosines of HHV-6B U69 and EBV BGLF4 corresponding to HSV-2 Tyr-162 on EF-1δ in cell cultures.** (a) COS-7 cells were transfected with a plasmid expressing EGFP-EF-1δ(F) (lanes 1-5) or a plasmid expressing EGFP-EF-1δ-S133A(F) (lane 6) combined with a plasmid expressing empty (lane 1), SE-U69 (lanes 2, 6), SE-U69-K219M (lane 3), SE-U69-Y207E (lane 4), or SE-U69-Y207F (lane 5), and harvested 48 h post-transfection. Cell lysates were analyzed by immunoblotting with antibodies to Flag-tag, EF-1δ-S133^P^, Strep-tag, or β-actin. Digital images are representative of three independent experiments. (b) Amount of EGFP-EF-1δ(F)-S133^P^ protein detected with anti-EF-1δ-S133^P^ monoclonal antibody (a, top panel) relative to that of EGFP-EF-1δ(F) protein detected with anti-Flag-tag antibody (a, second panel) in transfected cells. Data were normalized by dividing the sum of the data on the same blot^59^. Each value is the mean ± SEM of three experiments. Statistical significance was analyzed by ANOVA with the Tukey’s test. n.s., not significant. (c) COS-7 cells were transfected with a plasmid expressing EGFP-EF-1δ(F) (lanes 1-5) or a plasmid expressing EGFP-EF-1δ-S133A(F) (lane 6) and a plasmid expressing empty (lane 1), BGLF4 (lane 2, 6), BGLF4-K102I (lane 3), BGLF4-Y89E (lane 4), or BGLF4-Y89F (lane 5), and harvested 48 h post-transfection. Cell lysates were analyzed by immunoblotting with antibodies to Flag-tag, EF-1δ-S133^P^, BGLF4, or β-actin. Digital images are representative of nine independent experiments. (d) Amount of EGFP-EF-1δ(F)-S133^P^ protein detected with anti-EF-1δ-S133^P^ monoclonal antibody (a, top panel) relative to that of EGFP-EF-1δ(F) protein detected with anti-Flag-tag antibody (a, second panel) in transfected cells. Data were normalized by dividing the sum of the data on the same blot. Each value is the mean ± SEM of nine experiments. Statistical significance was analyzed by ANOVA with the Tukey’s test. n.s., not significant.

**S-Fig. 6. Effect of mutations in UL13 on viral replication and cell-cell spread in Vero cells. a, b.** Vero cells were infected with wild-type HSV-2 186, ΔUL13, ΔUL13-repair, UL13-K176M, UL13-K176M-repair, UL13-Y162E, UL13-Y162F, or UL13-Y162E/F-repair at an MOI of 0.01 (a) or 3 (b). Total virus titers in cell culture supernatants and infected cells were harvested at 24 h (a) or 12 h (b) post-infection and assayed. Each value is the mean ± SEM of four experiments. Statistical significance was analyzed by ANOVA with Tukey’s test. n.s., not significant. **c.** Vero cells were infected with wild-type HSV-2 186, ΔUL13, ΔUL13-repair, UL13-K176M, UL13-K176M-repair, UL13-Y162E, UL13-Y162F, or UL13-Y162E/F-repair at an MOI of 0.0001 under plaque assay conditions. Diameters of 20 single plaques for each virus were measured 48 h post-infection. Each data point is the mean ± SEM of the measured plaque sizes. Statistical significance was analyzed by ANOVA with Tukey’s test. n.s., not significant. Data are representative of three independent experiments.

**S-Fig. 7. Effects of mutations in UL13 Tyr-162 on mortality, viral replication, and pathogenic manifestation in vaginas of guinea pigs following intravaginal infection. a-c.** Eighteen 5-week-old female Hartley guinea pigs used in Fig. 4a, b were intravaginally infected with 1×10^4^ PFU UL13-Y162F or UL13-Y162E/F-repair. **a.** Survival of guinea pigs was monitored for 21 days post-infection. Statistical significance was analyzed by log-rank test. n.s., not significant. **b.** Vaginal secretions of guinea pigs at 3- and 5-days post-infection were harvested and virus titers were assayed. Dashed line indicates the limit of detection. Each data point is the virus titer of one guinea pig. Horizontal bars indicate the mean of each group. Statistical significance was analyzed by Mann-Whitney *U*-test. n.s., not significant. **c.** Clinical scores of guinea pigs at 21 days post-infection were monitored. Data are the mean of the observations. Statistical significance was analyzed by Mann-Whitney *U*-test. n.s., not significant.

**S-Fig. 8. Sequence alignment around the GxGxxG motif of host cellular PKs.** Sequence alignment around the GxGxxG motif of host cellular protein kinases. Amino acids of host cellular protein kinases are labeled with their NCBI Reference Sequence numbers. The 10 kinases of each major kinase group, excluding CDK, are shown. Highly conserved glycine residues in the GxGxxG motif and valine residue near the GxGxxG motif are in white. Tyrosine residues corresponding to CDK1 Tyr-15 are in pink.

**S-Fig. 9. Sequence alignment around the GxGxxG motif of other viral serine/threonine PKs.** Sequence alignment around the GxGxxG motif of F10L kinase homologs (a) and B1R kinase homologs (b) conserved in poxviruses, and that of Us3 kinase homologs conserved in the subfamily *Alphaherpesvirinae* (c). Tyrosines corresponding to CDK1 Tyr-15 are in pink. Amino acids of viral kinases are labeled with their NCBI gene identification numbers and virus names. Highly conserved glycine residues in the GxGxxG motif and valine residue near the GxGxxG motif are in white. VACV, vaccinia virus; MPV, monkeypox virus; VARV, variola virus; SwPV, swinepox virus; LSDV, lumpy skin disease virus; DPV, deerpox virus; SQPV, squirrel poxvirus; HSV-1, herpes simplex virus 1; HSV-2, herpes simplex virus 2; VZV, varicella-zoster virus; PRV, pseudorabies virus; MDV, Marek’s disease virus; SaHV-1, saimiriine herpesvirus 1; CaHV-1, canid herpesvirus 1; FeHV-1, feline herpesvirus 1; BoHV-1, bovine herpesvirus 1; CpHV-1, caprine herpesvirus 1; EHV-1, equine herpesvirus 1; EHV-4, equine herpesvirus 4.

## Notes

### Competing Interest Statement

The authors have declared no competing interest.

