## Supplementary material for "Regulatory Mimicry of Cyclin-Dependent Kinases by Conserved Herpesvirus Protein Kinases": S-Figure ans S-Table

**a**

GxGxxG motif

| Group | Accession | Virus | Protein | Start | | Sequence | | End |
|---|---|---|---|---|---|---|---|---|
| α | ADD59987.1 | HSV-1 | UL13 | 155 | : | S F G G S G G Y G - E V Q L I | : | 168 |
| α | BAF81492.1 | HSV-2 | UL13 | 155 | : | S F G G S G G Y G - E V Q L I | : | 168 |
| α | NP_040169.1 | VZV | ORF47 | 136 | : | R F A G R G T Y G - R V H I Y | : | 149 |
| α | AEM64098.1 | PRV | UL13 | 84 | : | V A V G A G S Y G - S V L V Y | : | 97 |
| α | AVI57767.1 | MDV | UL13 | 149 | : | I Y A G S G S Y G - V V K I F | : | 162 |
| α | YP_003933827.1 | SaHV-1 | UL13 | 224 | : | S F G G S G G Y G - E V E L L | : | 237 |
| α | YP_009252273.1 | CHV-1 | UL13 | 224 | : | V F G G A G G Y G - E V K I Y | : | 237 |
| α | AVW80427.1 | FeHV-1 | UL13 | 240 | : | K F G G A G G Y G - E V Q I Y | : | 253 |
| α | ALR87813.1 | BoHV-1 | UL13 | 119 | : | Q L R G A G G Y G - S V V V H | : | 132 |
| α | QBM10888.1 | CpHV-1 | UL13 | 111 | : | Q A R G A G S Y G - S V V V Y | : | 124 |
| α | P28966 | EHV-1 | UL13 | 227 | : | K F G G A G S Y G - E V Q I F | : | 240 |
| α | AAC59566.1 | EHV-4 | UL13 | 231 | : | K F G G A G S Y G - E V Q I F | : | 244 |
| β | P16788.1 | HCMV | UL97 | 335 | : | Y R L G Q G S F G - E V W P L | : | 348 |
| β | NP_042962.1 | HHV-6A | U69 | 199 | : | R V L G V G A Y G - K V F D L | : | 212 |
| β | AVK93842.1 | HHV-6B | U69 | 200 | : | R V L G V G A Y G - K V F D L | : | 213 |
| β | YP_073809.1 | HHV-7 | U69 | 183 | : | R I L G S G S Y G - M V Y D L | : | 196 |
| β | YP_214098.1 | MCMV | UL97 | 269 | : | T L I G K G S F G - Q V W R L | : | 282 |
| β | AUI40786.1 | RhCMV | Rh132 | 242 | : | R R I G C G S F G - E I W P I | : | 255 |
| γ | P13288.2 | EBV | BGLF4 | 82 | : | Y L L G R G S Y G - A V Y A H | : | 95 |
| γ | YP_001129389.1 | KSHV | ORF36 | 88 | : | K R L G R G A F G I I V P I S | : | 102 |
| γ | AAF19301.1 | MHV-68 | ORF36 | 85 | : | D T L G E G G F G - R V S T M | : | 98 |
| γ | NP_042633.1 | EHV-2 | ORF36 | 81 | : | L A L G K G E Y G - R V E A V | : | 94 |
| γ | YP_438159.1 | OvHV-2 | ORF36 | 81 | : | T L L A K G A F G - K V Y S S | : | 94 |

**b**

GxGxxG motif

| Accession | Species | Protein | Start | | Sequence | | End |
|---|---|---|---|---|---|---|---|
| NP_001777.1 | Human | CDK1 | 8 | : | E K I G E G T Y G - V V Y K G | : | 21 |
| NP_001789.2 | Human | CDK2 | 8 | : | E K I G E G T Y G - V V Y K A | : | 21 |
| NP_001249.1 | Human | CDK3 | 8 | : | E K I G E G T Y G - V V Y K A | : | 21 |
| NP_000066.1 | Human | CDK4 | 10 | : | A E I G V G A Y G - T V Y K A | : | 23 |
| NP_004926.1 | Human | CDK5 | 8 | : | E K I G E G T Y G - T V F K A | : | 21 |
| NP_001138778.1 | Human | CDK6 | 17 | : | A E I G E G A Y G - K V F K A | : | 30 |
| NP_001790.1 | Human | CDK7 | 16 | : | D F L G E G Q F A - T V Y K A | : | 29 |
| NP_001251.1 | Human | CDK8 | 25 | : | C K V G R G T Y G - H V Y K A | : | 38 |
| NP_001252.1 | Human | CDK9 | 23 | : | A K I G Q G T F G - E V F K A | : | 36 |
| NP_443714.3 | Human | CDK10 | 43 | : | N R I G E G T Y G - I V Y R A | : | 56 |
| NP_076916.2 | Human | CDK11A | 427 | : | N R I E E G T Y G - V V Y R A | : | 440 |
| NP_001778.2 | Human | CDK11B | 442 | : | N R I E E G T Y G - V V Y R A | : | 455 |
| NP_057591.2 | Human | CDK12 | 731 | : | G I I G E G T Y G - Q V Y K A | : | 744 |
| NP_003709.3 | Human | CDK13 | 709 | : | G I I G E G T Y G - Q V Y K A | : | 722 |
| NP_001274064.1 | Human | CDK14 | 139 | : | E K L G E G S Y A - T V Y K G | : | 152 |
| NP_001248364.1 | Human | CDK15 | 107 | : | E K L G E G S Y A - T V Y K G | : | 120 |
| NP_006192.1 | Human | CDK16 | 169 | : | D K L G E G T Y A - T V Y K G | : | 182 |
| NP_002586.2 | Human | CDK17 | 196 | : | E K L G E G T Y A - T V Y K G | : | 209 |
| NP_997668.1 | Human | CDK18 | 178 | : | D K L G E G T Y A - T V F K G | : | 191 |
| NP_055891.1 | Human | CDK19 | 25 | : | C K V G R G T Y G - H V Y K A | : | 38 |
| NP_848519.1 | Human | CDK20 | 8 | : | G R I G E G A H G - I V F K A | : | 21 |

**N. Koyanagi et al. S-Fig. 1**

**S-Fig. 1. Sequence alignment around the GxGxxG motif of CHPKs and human CDKs. a.** CHPKs amino acids are labeled with their NCBI gene identification numbers and virus names. HSV-1, herpes simplex virus 1; HSV-2, herpes simplex virus 2; VZV, varicella-zoster virus; PRV, pseudorabies virus; MDV, Marek's disease virus; SaHV-1, saimiriine herpesvirus 1; CaHV-1, canid herpesvirus 1; FeHV-1, feline herpesvirus 1; BoHV-1, bovine herpesvirus 1; CpHV-1, caprine herpesvirus 1; EHV-1, equine herpesvirus 1; EHV-4, equine herpesvirus 4; HCMV, human cytomegalovirus; HHV-6A, human herpesvirus 6A; HHV-6B, human herpesvirus 6B; HHV-7, human herpesvirus 7; MCMV, murine cytomegalovirus; RhCMV, rhesus cytomegalovirus; EBV, Epstein-Barr virus; KSHV, Kaposi's sarcoma-associated herpesvirus; MHV-68, murine gammaherpesvirus-68; EHV-2, equine herpesvirus 2; and OvHV-2, ovine herpesvirus 2. HSV-1, HSV-2, VZV, PRV, MDV, SaHV-1, CaHV-1, FeHV-1, BoHV-1, CpHV-1, EHV-1 and EHV-4 belong to the subfamily *Alphaherpesvirinae*; HCMV, HHV-6A, HHV-6B, HHV-7, MCMV and RhCMV belong to the *Betaherpesvirinae*; and EBV, KSHV, MHV-68, EHV-2 and OvHV-2 belong to the *Gammaherpesvirinae*. Highly conserved glycine residues in the GxGxxG motif and valine residue near the GxGxxG motif are in white. Tyrosine residues corresponding to the CDK1 Tyr-15 position of the GxGxxG motif are in pink. **b.** CDK amino acids are labeled with their NCBI gene identification numbers. Highly conserved glycine residues in the GxGxxG motif and valine residue near the GxGxxG motif are in white. Tyrosine residues corresponding to the CDK1 Tyr-15 position of the GxGxxG motif are in pink.

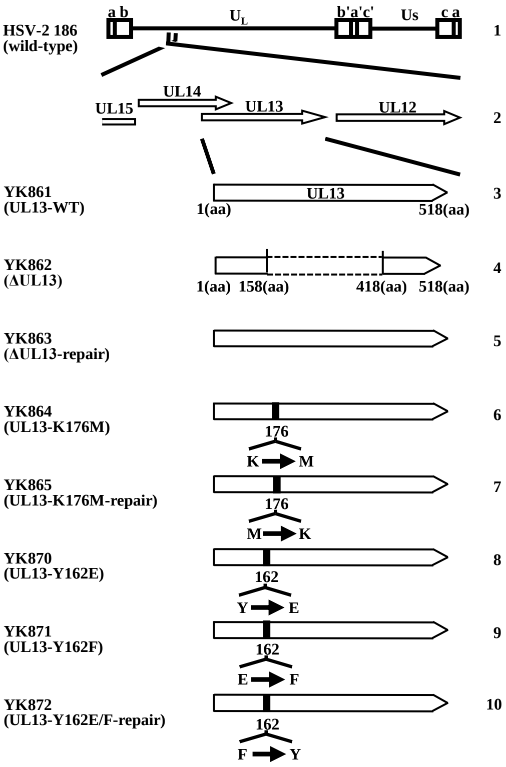


**S-Fig. 2. Schematic diagrams of the genomic structures of wild-type HSV-2 186 and the relevant domains of recombinant viruses used in this study.** Line 1, wild-type HSV-2 186 genome; Line 2, domain of the UL12 gene to the UL15 gene; Lines 3 to 10, recombinant viruses with mutations in the UL13 gene.

**N. Koyanagi et al. S-Fig. 2**

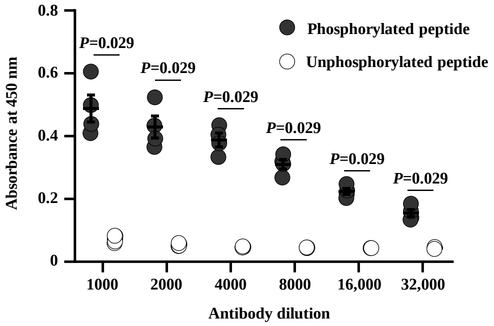


**S-Fig. 3. Generation of rabbit polyclonal antibodies to UL13-Y162$^{P}$**. ELISA was performed to assess the specificity of UL13-Y162$^{P}$ polyclonal antibodies. Phosphorylated Tyr-162 peptide of UL13 (Cys-GGSGG(pY)GEVQL) or non-phosphorylated Tyr-162 peptide of UL13 (Cys-GGSGGYGEVQL) was used for ELISAs. Each value is the mean ± SEM of four experiments. Statistical significance was analyzed by Mann-Whitney *U*-test.

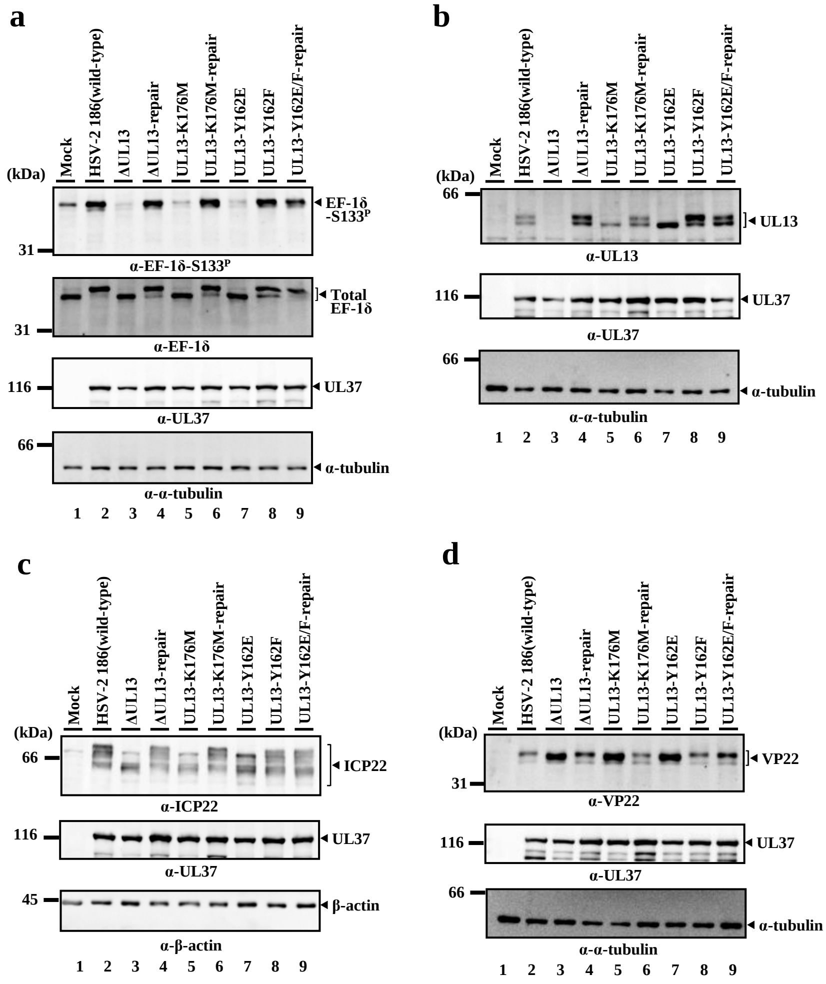


**S-Fig. 4. Effects of mutations in UL13 on post-translational processing of UL13 substrates in Vero cells.** Vero cells mock-infected or infected with wild-type HSV-2 186, ΔUL13, ΔUL13-repair, UL13-K176M, UL13-K176M-repair, UL13-Y162E, UL13-Y162F, or UL13-Y162E/F-repair for 24 h at an MOI of 3 were analyzed by immunoblotting with antibodies to EF-1δ (a), EF-1δ-S133$^{P}$ (a), UL13 (b), ICP22 (c), VP22 (d), UL37 (a-d), α-tubulin (a, b, d), or β-actin (c). Digital images are representative of three independent experiments.

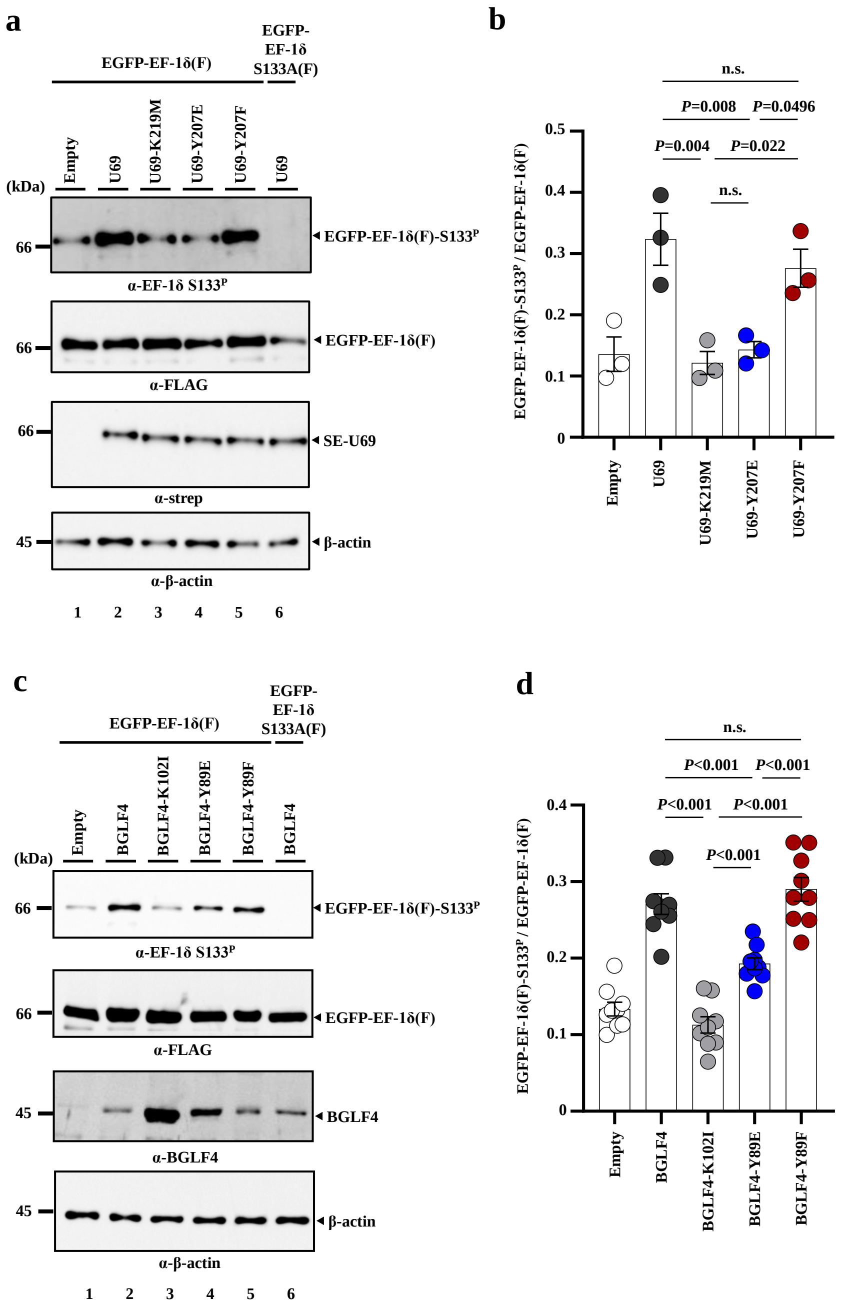


N. Koyanagi et al. S-Fig. 5

**S-Fig. 5. Effects of mutations in tyrosines of HHV-6B U69 and EBV BGLF4 corresponding to HSV-2 Tyr-162 on EF-1δ in cell cultures.** (a) COS-7 cells were transfected with a plasmid expressing EGFP-EF-1δ(F) (lanes 1-5) or a plasmid expressing EGFP-EF-1δ-S133A(F) (lane 6) combined with a plasmid expressing empty (lane 1), SE-U69 (lanes 2, 6), SE-U69-K219M (lane 3), SE-U69-Y207E (lane 4), or SE-U69-Y207F (lane 5), and harvested 48 h post-transfection. Cell lysates were analyzed by immunoblotting with antibodies to Flag-tag, EF-1δ-S133$^{P}$, Strep-tag, or β-actin. Digital images are representative of three independent experiments. (b) Amount of EGFP-EF-1δ(F)-S133$^{P}$ protein detected with anti-EF-1δ-S133$^{P}$ monoclonal antibody (a, top panel) relative to that of EGFP-EF-1δ(F) protein detected with anti-Flag-tag antibody (a, second panel) in transfected cells. Data were normalized by dividing the sum of the data on the same blot[59]. Each value is the mean ± SEM of three experiments. Statistical significance was analyzed by ANOVA with the Tukey's test. n.s., not significant. (c) COS-7 cells were transfected with a plasmid expressing EGFP-EF-1δ(F) (lanes 1-5) or a plasmid expressing EGFP-EF-1δ-S133A(F) (lane 6) and a plasmid expressing empty (lane 1), BGLF4 (lane 2, 6), BGLF4-K102I (lane 3), BGLF4-Y89E (lane 4), or BGLF4-Y89F (lane 5), and harvested 48 h post-transfection. Cell lysates were analyzed by immunoblotting with antibodies to Flag-tag, EF-1δ-S133$^{P}$, BGLF4, or β-actin. Digital images are representative of nine independent experiments. (d) Amount of EGFP-EF-1δ(F)-S133$^{P}$ protein detected with anti-EF-1δ-S133$^{P}$ monoclonal antibody (a, top panel) relative to that of EGFP-EF-1δ(F) protein detected with anti-Flag-tag antibody (a, second panel) in transfected cells. Data were normalized by dividing the sum of the data on the same blot. Each value is the mean ± SEM of nine experiments. Statistical significance was analyzed by ANOVA with the Tukey's test. n.s., not significant.

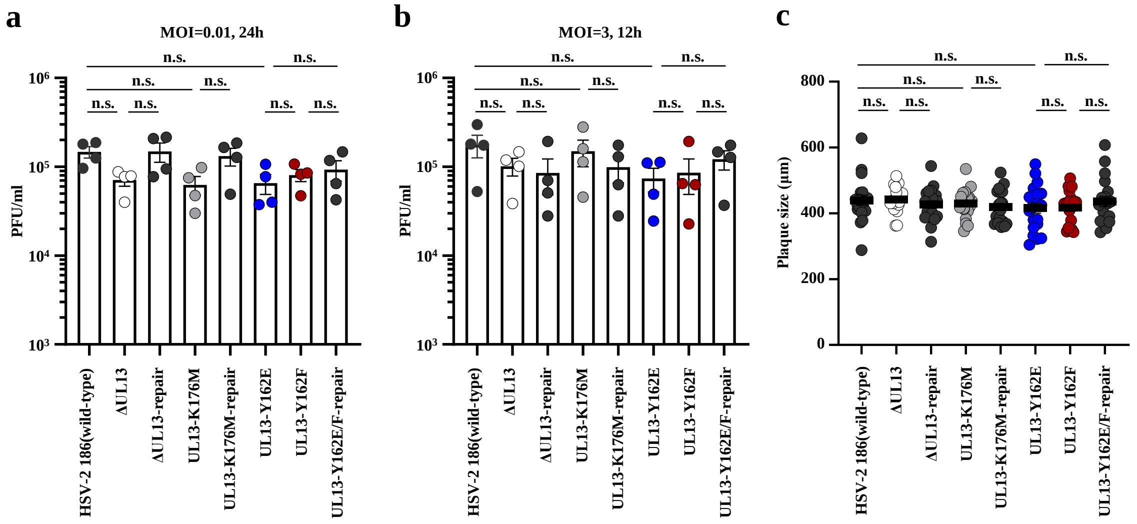


**S-Fig. 6. Effect of mutations in UL13 on viral replication and cell-cell spread in Vero cells. a, b.** Vero cells were infected with wild-type HSV-2 186, ΔUL13, ΔUL13-repair, UL13-K176M, UL13-K176M-repair, UL13-Y162E, UL13-Y162F, or UL13-Y162E/F-repair at an MOI of 0.01 (a) or 3 (b). Total virus titers in cell culture supernatants and infected cells were harvested at 24 h (a) or 12 h (b) post-infection and assayed. Each value is the mean ± SEM of four experiments. Statistical significance was analyzed by ANOVA with Tukey's test. n.s., not significant. **c.** Vero cells were infected with wild-type HSV-2 186, ΔUL13, ΔUL13-repair, UL13-K176M, UL13-K176M-repair, UL13-Y162E, UL13-Y162F, or UL13-Y162E/F-repair at an MOI of 0.0001 under plaque assay conditions. Diameters of 20 single plaques for each virus were measured 48 h post-infection. Each data point is the mean ± SEM of the measured plaque sizes. Statistical significance was analyzed by ANOVA with Tukey's test. n.s., not significant. Data are representative of three independent experiments.

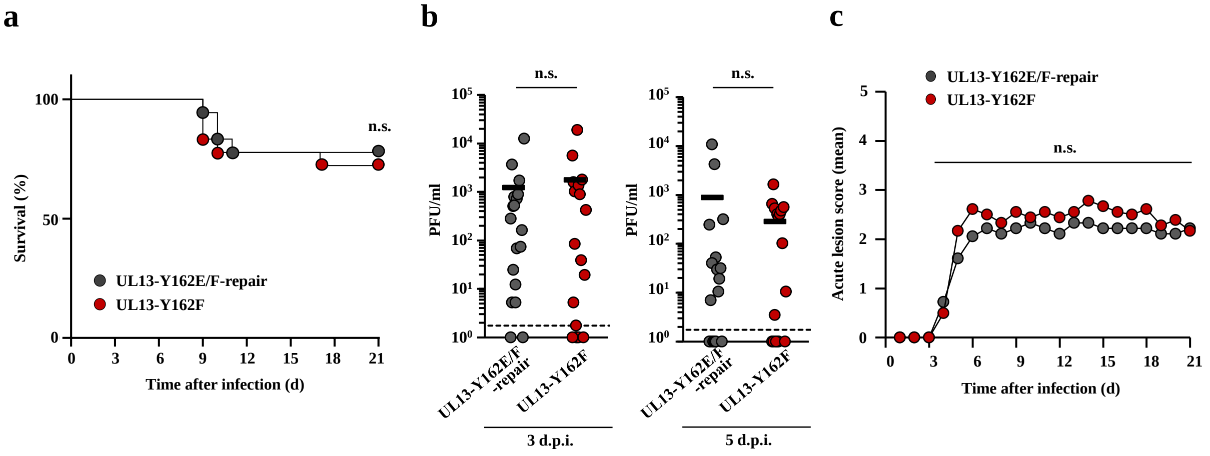


**S-Fig. 7. Effects of mutations in UL13 Tyr-162 on mortality, viral replication, and pathogenic manifestation in vaginas of guinea pigs following intravaginal infection. a-c.** Eighteen 5-week-old female Hartley guinea pigs used in Fig. 4a, b were intravaginally infected with $1\times10^4$ PFU UL13-Y162F or UL13-Y162E/F-repair. **a.** Survival of guinea pigs was monitored for 21 days post-infection. Statistical significance was analyzed by log-rank test. n.s., not significant. **b.** Vaginal secretions of guinea pigs at 3- and 5-days post-infection were harvested and virus titers were assayed. Dashed line indicates the limit of detection. Each data point is the virus titer of one guinea pig. Horizontal bars indicate the mean of each group. Statistical significance was analyzed by Mann-Whitney *U*-test. n.s., not significant. **c.** Clinical scores of guinea pigs at 21 days post-infection were monitored. Data are the mean of the observations. Statistical significance was analyzed by Mann-Whitney *U*-test. n.s., not significant.

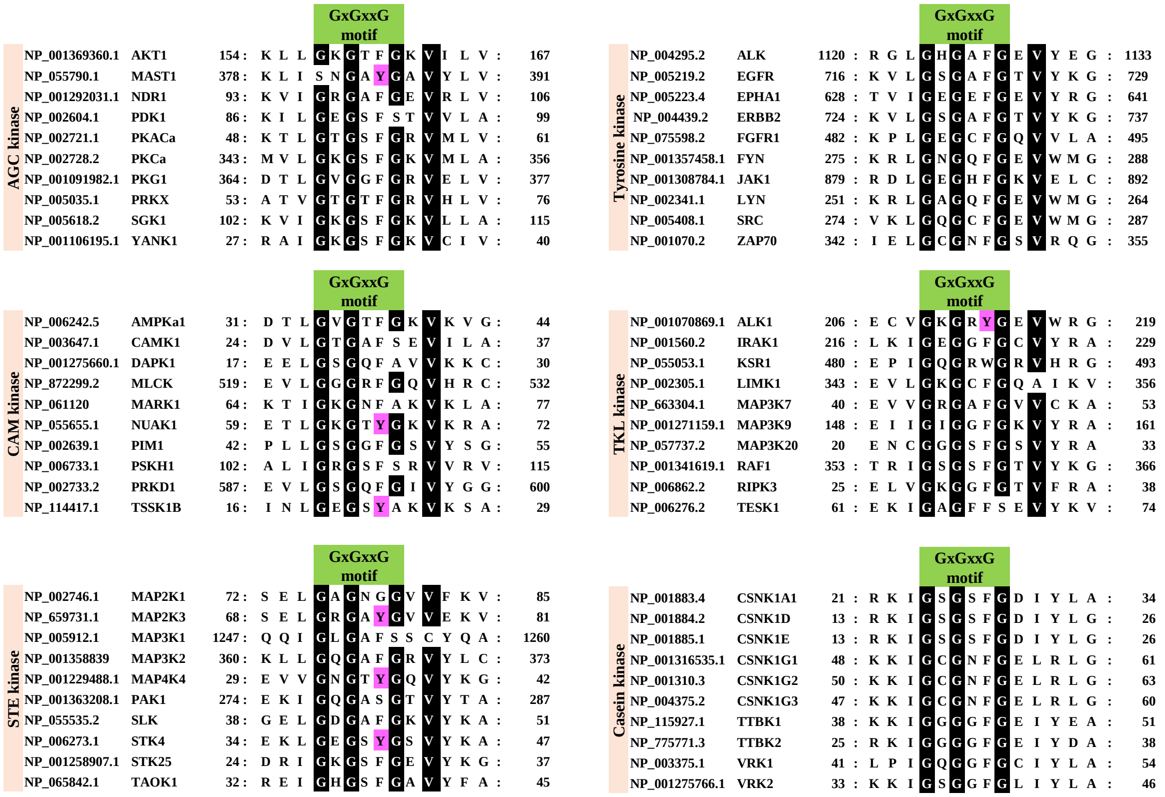


**S-Fig. 8. Sequence alignment around the GxGxxG motif of host cellular PKs.** Sequence alignment around the GxGxxG motif of host cellular protein kinases. Amino acids of host cellular protein kinases are labeled with their NCBI Reference Sequence numbers. The 10 kinases of each major kinase group, excluding CDK, are shown. Highly conserved glycine residues in the GxGxxG motif and valine residue near the GxGxxG motif are in white. Tyrosine residues corresponding to CDK1 Tyr-15 are in pink.

**N. Koyanagi et al. S-Fig. 8**

**a**

| | | | | GxGxxG motif | |
|---|---|---|---|---|---|
| YP_232931.1 | VACV | F10L | 91 | : Y H I S T G G Y G - I V F K I : | 104 |
| ADK39065.1 | MPV | C16L | 91 | : Y H I S T G G Y G - I V F K I : | 104 |
| DAA80539.1 | VARV | C14L | 91 | : Y H I S T G G Y G - I V F K I : | 104 |
| NP_570182.1 | SwPV | C20L | 89 | : Y H I S T G G Y G - I V F K M : | 102 |
| NP_150459.1 | LSDV | F10L | 93 | : Y H I S T G G Y G - I V F K I : | 106 |
| YP_227408.1 | DPV | F10L | 92 | : Y H I S T G G Y G - I V F K I : | 105 |
| YP_008658436.1 | SQPV | F10L | 90 | : Y H L S T G G Y G - I V F R L : | 103 |

**b**

| | | | | GxGxxG motif | |
|---|---|---|---|---|---|
| YP_233065.1 | VACV | B1R | 20 | : P L I G K G G F G - S I Y T T : | 33 |
| ADK39065.1 | MPV | B3R | 20 | : P L I G K G G F G - S I Y T T : | 33 |
| DAA80679.1 | VARV | B1R | 20 | : P L I G K G G F G - S I Y T T : | 33 |
| NP_570297.1 | SPV | B1R | 26 | : T L I G K G G F G - C I Y T A : | 39 |
| NP_150573.1 | LSDV | B1R | 27 | : K I I G Q G G F G - F I F L A : | 40 |
| YP_227529.1 | DPV | B1R | 28 | : N I I G K G G F G - F I Y L S : | 41 |

**c**

| | | | | GxGxxG motif | |
|---|---|---|---|---|---|
| ADD60050.1 | HSV-1 | Us3 | 195 | : G A L T P G S E G - C V F D S : | 208 |
| YP009137215.1 | HSV-2 | Us3 | 195 | : G A L I P G S E G - C V F D S : | 208 |
| CAA27949.1 | VZV | ORF66 | 97 | : K T F T P G A E G - F A F A C : | 110 |
| AEM64106.1 | PRV | Us3 | 59 | : Q P L Q S G S E G - R V F V A : | 72 |
| AVI57869.1 | MDV | Us3 | 106 | : S S L P P G S E G - Y I Y V C : | 119 |
| YP_003933845.1 | SaHV-1 | Us3 | 159 | : D T L T P G S E G - C V F V A : | 172 |
| YP_009252293.1 | CaHV-1 | Us3 | 113 | : K S L T P G S E G - R V F I A : | 126 |
| AVW80446.1 | FeHV-1 | Us3 | 68 | : K T L T P G S E G - R V F V A : | 81 |
| ALR87834.1 | BoHV-1 | Us3 | 169 | : R R L T P G S E G - R V F E A : | 182 |
| QBM10906.1 | CpHV-1 | Us3 | 99 | : E T L T P G S E G - R V F V A : | 112 |
| BAX36465.1 | EHV-1 | Us3 | 97 | : K T L T P G S E G - R V M V A : | 110 |
| AAC59588.1 | EHV-4 | Us3 | 99 | : K T L T P G S E G - R V M V A : | 112 |

**S-Fig. 9. Sequence alignment around the GxGxxG motif of other viral serine/threonine PKs.** Sequence alignment around the GxGxxG motif of F10L kinase homologs (a) and B1R kinase homologs (b) conserved in poxviruses, and that of Us3 kinase homologs conserved in the subfamily *Alphaherpesvirinae* (c). Tyrosines corresponding to CDK1 Tyr-15 are in pink. Amino acids of viral kinases are labeled with their NCBI gene identification numbers and virus names. Highly conserved glycine residues in the GxGxxG motif and valine residue near the GxGxxG motif are in white. VACV, vaccinia virus; MPV, monkeypox virus; VARV, variola virus; SwPV, swinepox virus; LSDV, lumpy skin disease virus; DPV, deerpox virus; SQPV, squirrel poxvirus; HSV-1, herpes simplex virus 1; HSV-2, herpes simplex virus 2; VZV, varicella-zoster virus; PRV, pseudorabies virus; MDV, Marek's disease virus; SaHV-1, saimiriine herpesvirus 1; CaHV-1, canid herpesvirus 1; FeHV-1, feline herpesvirus 1; BoHV-1, bovine herpesvirus 1; CpHV-1, caprine herpesvirus 1; EHV-1, equine herpesvirus 1; EHV-4, equine herpesvirus 4.

S-Table 1. Oligonucleotide sequences for the construction of plasmids.

| Plasmid | Sequence (5'-3') |
|---|---|
| pGEX-ICP22-P1 | GCGAATTCATGGCAGACATCCCCCCGGA |
| | GCGTCGACCTAGATAGGTCTGTCGCTTCTCC |
| pME-BGLF4-Y89E | CTGGGGCGGGGGAGCGAAGGGGCCGTGTATGCA |
| | TGCATACACGGCCCCTTCGCTCCCCCGCCCCAG |
| pME-BGLF4-Y89F | CTGGGGCGGGGGAGCTTCGGGGCCGTGTATGCA |
| | TGCATACACGGCCCCGAAGCTCCCCCGCCCCAG |
| pcDNA-SE-U69 | ATCGAATTCATGGACAACGGTGTGGAGACACCTCAAGGTC |
| | TATGCGGCCGCTCACATCTGAAAGAGAGATGATG |
| pcDNA-SE-U69-K219M | GATAAAGTGGCCATAATGACGGCCAACGAAGAC |
| | GTCTTCGTTGGCCGTCATTATGGCCACTTTATC |
| pcDNA-SE-U69-Y207E | CTCGGAGTGGGTGCCGAGGGGAAGGTGTTTGAT |
| | ATCAAACACCTTCCCCTCGGCACCCACTCCGAG |
| pcDNA-SE-U69-Y207F | CTCGGAGTGGGTGCCTTCGGGAAGGTGTTTGAT |
| | ATCAAACACCTTCCCGAAGGCACCCACTCCGAG |

S-Table 2. Oligonucleotide sequences for the construction of recombinant viruses.

| Recombinant virus | Sequence (5’-3’) |
|---|---|
| UL13-Y162E | ATCCCCGGGGCCCGCAGCTTCGGGGGCTCGGGGGGGGAGGGCGAGGTGCAGTTGATTCGAGGATGACGACGATAAGTAGGG |
| | TCACGGCGAGCTTGTGTTCGCGAATCAACTGCACCTCGCCCTCCCCCCCCGAGCCCCCGAAGCCAACCAATTAACCAATTCTGATTAG |
| UL13-Y162F | ATCCCCGGGGCCCGCAGCTTCGGGGGCTCGGGGGGGTTCGGCGAGGTGCAGTTGATTCGAGGATGACGACGATAAGTAGGG |
| | TCACGGCGAGCTTGTGTTCGCGAATCAACTGCACCTCGCCGAACCCCCCCGAGCCCCCGAAGCCAACCAATTAACCAATTCTGATTAG |
| UL13-Y162E/F-repair | ATCCCCGGGGCCCGCAGCTTCGGGGGCTCGGGGGGGTACGGCGAGGTGCAGTTGATTCGAGGATGACGACGATAAGTAGGG |
| | TCACGGCGAGCTTGTGTTCGCGAATCAACTGCACCTCGCCGTACCCCCCCGAGCCCCCGAAGCCAACCAATTAACCAATTCTGATTAG |
